# SiamEEGNet: Siamese Neural Network-Based EEG Decoding for Drowsiness Detection

**DOI:** 10.1101/2023.10.23.563513

**Authors:** Li-Jen Chang, Hsi-An Chen, Chin Chang, Chun-Shu Wei

**Affiliations:** Department of Computer Science, National Yang Ming Chiao Tung University (NYCU), Hsinchu, Taiwan; Institute of Education, the Institute of Biomedical Engineering, and the Brain Science and Technology Center, NYCU, Hsinchu, Taiwan

**Keywords:** EEG, Drowsiness detection, Siamese network

## Abstract

Recent advancements in deep-learning have significantly enhanced EEG-based drowsiness detection. However, most existing methods overlook the importance of relative changes in EEG signals compared to a baseline, a fundamental aspect in conventional EEG analysis including event-related potential and time-frequency spectrograms. We herein introduce SiamEEGNet, a Siamese neural network architecture designed to capture relative changes between EEG data from the baseline and a time window of interest. Our results demonstrate that SiamEEGNet is capable of robustly learning from high-variability data across multiple sessions/subjects and outperforms existing model architectures in cross-subject scenarios. Furthermore, the model’s interpretability associates with previous findings of drowsiness-related EEG correlates. The promising performance of SiamEEGNet highlights its potential for practical applications in EEG-based drowsiness detection. We have made the source codes available at http://github.com/CECNL/SiamEEGNet.

## I. Introduction

Drowsy driving is a major contributor to traffic accidents and results in significant human and financial costs. In the United States, approximately 15-33 percent of fatal crashes are associated with drowsy driving [1]. Drowsiness can be caused by various factors, such as sleepiness, changes in circadian rhythm due to shift work, sleep deprivation, and fatigue, all of which greatly affect alertness, concentration, and reaction time [2]. To minimize the negative effects of drowsy driving and detect drowsiness in its early stages, it is essential to monitor drowsiness. Generally, research on drowsiness detection can be categorized into three groups based on the modalities used to monitor drowsiness: vehicle-based, behavioral-based, and physiological-based [3]. Physiological-based methods, such as electroencephalography (EEG), have the advantage of directly measuring and reflecting brain activity associated with drowsiness in the driver [4]. Thus, they are considered the most promising approach for detecting drowsiness. EEG is a widely used tool to monitor and analyze brain activity, and many studies have shown that it is feasible to use EEG to detect drowsiness [5], [6]. However, the EEG signal is characterized by being non-stationary and can vary greatly between subjects, as well as within the same subject [7], [8]. This variability requires careful analysis and interpretation to accurately understand and explore the human brain. Due to the time-resolved nature of the EEG signal, it is susceptible to temporal drift, which is not directly related to the experiment. Both physical factors, such as changes in sensor impedance, and mental factors, such as changes in mental condition, can contribute to this variability [9]. This phenomenon can have a significant impact on the analysis of EEG, as the baseline level of EEG activity changes over time.

With recent advances in deep learning (DL), an increasing number of researchers have focused on DL-based EEG decoding. For example, convolutional neural networks (CNNs) have been widely adopted for EEG classification tasks, due to their ability to extract high-level temporal and spatial information from raw EEG signals. CNN-based EEG decoding models transform raw EEG signals into a latent space and use learnable convolution kernels to extract information [10]–[12]. The convolutional kernels can achieve the effects of temporal and spatial filtering, akin to conventional approaches but without the constraints of predefined or hand-crafted features. The DL approach offers new opportunities to explore and exploit unconventional or high-level information in EEG dynamics and thus facilitates the understanding of brain dynamics.

To enhance performance of decoding drowsiness-related brain activities, many researchers have incorporated DL models into their studies. Four main DL models are used in most studies: CNN [13], [14], RNN [15], [16], Transformer [17], [18], and GCN [19], [20]. Although DL-based models have demonstrated significant improvements in decoding drowsy brain activities, most of them do not consider the relative change of brain activities, which is a crucial aspect when dealing with EEG signals. As mentioned earlier, the inter- and intra-subject variability in EEG poses a significant obstacle in performing cross-subject tasks. Relative change can be a potential solution to alleviate this problem, and several conventional EEG analysis approaches have already adopted this concept. One commonly used method is the baseline correction in the analysis of event-related potential (ERP) [21]. The approach involves subtracting the average voltage value of the EEG activity during the baseline period, which is the time prior to an external stimulation, from the EEG activity during the post-stimulus interval, which is the time following the stimulation. This helps correct for temporal drift in EEG signals. In the field of alertness/drowsiness monitoring, it has been observed that the brain dynamics associated with drowsiness can vary greatly between individuals, and even for the same individual, the level of alertness or drowsiness can vary at different times [22]. To address this issue and improve generalizability, many studies in this field take the relative power or the power ratio into consideration. This approach helps reduce individual differences and enhance the performance of alertness/drowsiness estimation [23], [24]. Moreover, there are numerous applications for EEG analysis and brain-computer interface (BCI) that make use of the relative change in EEG signals or its power [25]–[27]. These examples suggest that the exact voltage value or power of EEG signals is often not a sufficient criterion for EEG analysis. Therefore, it is common to employ a baseline or reference, typically in the form of the mean voltage or power of the signal, to represent the initial level for each subject or trial. As a result, the focus of many EEG analysis is on the relative change in brain activity, as this is believed to improve the robustness and generalizability for EEG analysis.

However, there are still some obstacles to overcome if we want to utilize the advantage of using the relative change in the EEG pattern in DL-based methods. For conventional single-branch CNN-based EEG decoding models, they are constrained by their structures that extracting information from a single input. This makes it difficult to capture the pattern of relative relationships or changes between trials. RNN-based models can capture short-term relative changes, but struggle with long-term considerations. As for the transformer-based model, it has tremendous potential to capture various features of EEG due to its attention mechanism. However, the biggest barrier to applying the transformer to EEG decoding is the small scale of EEG datasets. Neglecting the consideration of relative change in EEG can result in suboptimal performance in the scenario of cross-subject training or the requirement of additional cross-subject training techniques [28], [29]. Therefore, to utilize the advantage of using relative changes in the EEG pattern in DL-based methods, it is necessary to develop an alternative architecture, as shown in Fig. 1, that excels at learning changes between trials and extract different EEG patterns of relative change.

**Fig. 1:**
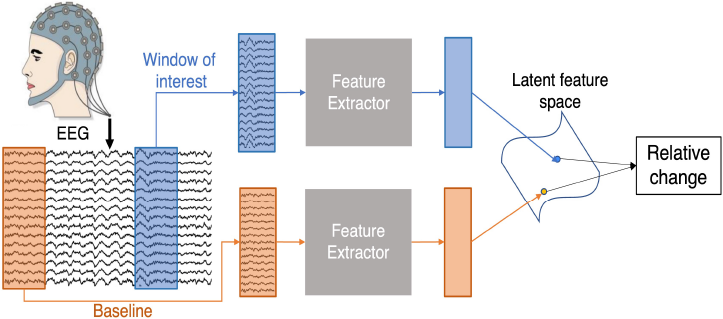
A Siamese network structure facilitates the extraction of features that represent the relative change between an EEG segment within a time window of interest and a baseline segment.

In this study, our primary objective is to enhance the performance of the drowsiness estimation in a DL-based EEG decoding model by introducing the concept of relative change. Therefore, we propose a novel Siamese neural network for EEG decoding, SiamEEGNet, to estimate drowsiness levels. The model is inspired by the Siamese network architecture [30] and incorporates modules adapted from novel DL-base EEG decoders. The Siamese architecture enables the extraction of consistent latent representations from two parallel inputs, and the shared feature extractor facilitates the learning of relative relationships between two inputs in the latent domain. This property enables learning and inference regarding the relative change in both the input data and the output labels. Thus, we can estimate the subject’s drowsy level by comparing the current trial with the baseline trial, which represents the level of alert baseline for each subject. Our key contributions are threefold. First, we present a novel Siamese EEG decoding architecture designed to capture the EEG pattern of relative change. This architecture is applied to the problem of drowsy EEG decoding and demonstrates outstanding performance in both the within-subject and cross-subject scenarios compared to existing methods. Second, we develop various manipulation techniques to improve stability and robustness in drowsy EEG decoding. Lastly, we extract visual evidence from SiamEEGNet to interpret the significant characteristics with neuroscientific insights.

## II. RelatedWork

### A. EEG-based drowsiness estimation

In recent years, due to the utilization of deep learning, significant advancements have been made in the field of EEG-based drowsiness estimation through the utilization of deep learning. Conventional machine learning approaches typically involve a two-step process, which includes extracting handcrafted features from EEG data and a prediction algorithm. Compared to machine learning approaches, DL-based methods enable an end-to-end training approach, allowing for datadriven feature extraction directly from EEG. Recent studies on DL-based drowsiness estimation can be broadly categorized into four types: CNN-based, RNN-based, Transformer-based and GCN-based methods. CNN-based models excel in extracting both temporal and spatial features through various convolutional kernel designs [13], [14]. Due to the basic characteristic of EEG (i.e., a time sequence), [15], [16] integrate the RNN-based method into their framework to take temporal dependencies into consideration. Transformer-based methods, known for their efficacy in handling sequential information, have shown promising results in various research fields [31]. By leveraging attention mechanisms, transformer-based methods can achieve performance comparable or even superior to RNN-based approaches [32]. Consequently, researchers have started exploring the application of Transformer-based methods in drowsiness detection, aiming to overcome the limitations associated with long-term sequences often encountered by RNN-based methods [17], [18]. Furthermore, the graph convolutional network (GCN) has become a popular choice for EEG-based drowsiness detection [33]. GCN focuses on learning spatial dependencies among EEG channels by treating multichannel EEG signals as graph data. Each EEG channel is represented as a node, and the relationships between two channels are captured by edges in the graph structure [19], [20].

### B. Siamese Network

A Siamese network is a neural network architecture comprising two identical sub-networks joined at their outputs. It was initially proposed for signature verification [30]. Later, the Siamese network architecture became widely adopted for similarity learning with deep neural networks. In most applications, the Siamese network applies the same transformation to both inputs and computes the similarity (or distance) between the latent representations of these inputs. For instance, it has been effectively used in tasks such as face verification [34] and object tracking, where the objective is to compare an exemplar image with candidate images and return a score representing the similarity of the two inputs [35].

Recently, the Siamese architecture has found applications in the field of EEG-based brain-computer interfaces (BCI). Many studies have utilized this architecture for similarity learning by minimizing the distance within samples of the same class while maximizing the distance between samples from different classes. By employing a Siamese network, it becomes possible to minimize the distance between samples of the same class, ensuring greater separation between samples from different classes. In this context, S. Zhang, et al. [36] used the Siamese network to learn distance-based representations from pairwise EEG data. S. Shahtalebi, et al. [37] developed a Siamese network and adopted a binary classification strategy for motor imagery classification tasks. Additionally, Yao Li, et al. [38] and X. Zhang, et al. [39] introduced convolutional correlation analysis (Conv-CA) and bidirectional Siamese correlation analysis, respectively, to enhance the performance of frequency classification of steady-state visual evoked potentials (SSVEP). These methods aim to compare EEG signals and reference signals in their latent representations and calculate the correlation between the two inputs.

## III. Architecture

In this section, we present the architecture of the SiamEEGNet, as shown in Fig.2. The proposed architecture consists of two identical feature extraction (FE) modules with identical model parameters. The objective is to map two inputs to a common latent space, which enables the prediction of the relative change in the drowsiness index (DI) using a regression layer. In the subsequent subsection, we provide a detailed explanation of each component in the SiamEEGNet, including the feature extraction module, multi-window processing, smoothing layer, and regression layer.

**Fig. 2:**
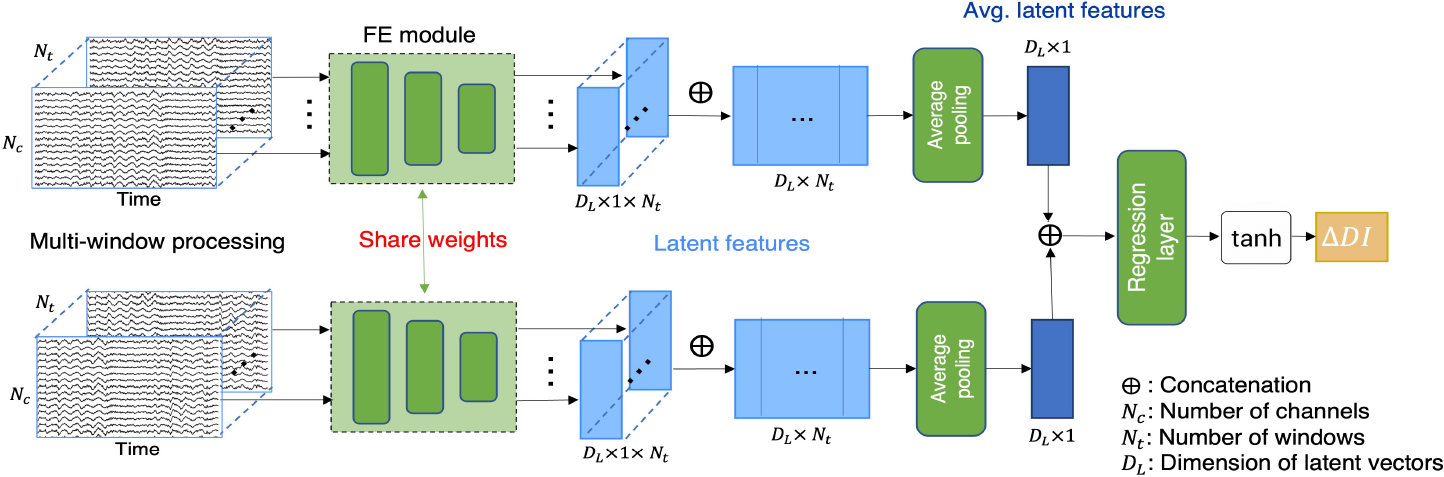
The proposed SiamEEGNet consists of two sub-networks that share the same structure and parameters. Each subnetwork comprises a feature extraction (FE) module and an average pooling layer. Additionally, a regression head is attached to both sub-networks to integrate information and make predictions based on the corresponding relative change in drowsiness level between the two inputs.

**Fig. 3:**
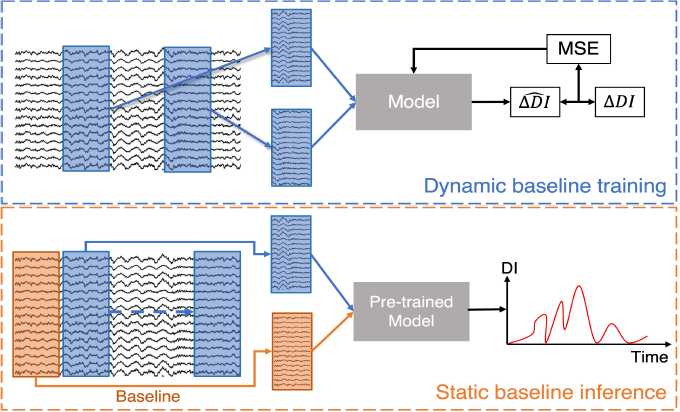
An illustration of the proposed dynamic baseline training technique and the static baseline inference. During the training stage, dynamic baseline training enables learning the relative change between any two segments within a session. In the inference stage, a static baseline is applied across the entire inference phase.

### A. Feature extraction module

The primary objective of the feature extraction module is to derive features conducive to the decoding of brain activities. To facilitate feature extraction from EEG data, SiamEEGNet requires a feature extraction module capable of capturing EEG data characteristics. As the default configuration, we integrate the feature extraction module from the established EEG decoding model, EEGNet [11], which consists of three main blocks: two convolutional blocks (temporal and depthwise separable convolution) and the classification block. The core aim of this module is to extract features from EEG signals and map them to a common latent space. Thus, we retain only the first two blocks of EEGNet for our feature extraction module. Moreover, the feature extraction module can be replaced with different EEG decoding models.

### B. Multi-window processing

To mitigate the short-term impact in the EEG recording, we developed a multi-window processing approach. The fundamental concept is to analyze a relatively extended EEG signal to identify a consistent brain state. Multi-window processing stacked *N*_*t*_ trials prior to the current trial together, where *N*_*t*_ is set to 10 as this approximately covers the trials within 90-second causal window. The *N*_*t*_ trials are then fed into the feature extractors individually, yielding the corresponding latent features. The calculation of feature extraction module incorporating multi-window processing can be presented as follows:

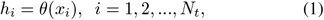

where *θ* refers to the feature extraction module, and *x*_*i*_ represents the *i*^*th*^ input trial in the multi-window processing. *h*_*i*_ is the corresponding latent feature with the size of (*D*_*L*_, 1), where *D*_*L*_ denotes the dimension of the latent features. In this study, we set *N*_*t*_ = 10 when using EEGNet as the feature extraction module.

### C. Regression layer

To predict the difference between two average latent features, we concatenate two outputs generated by the two parallel networks and apply a regression layer to predict the difference in DI between two input trials (ΔDI). The hyperbolic tangent was used as the output activation function to constrain the range of ΔDI within -1 to 1 (the range of ΔDI). The operation of the regression layer is shown as follows:

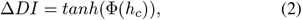

where *h*_*c*_ is the feature vector concatenating two average power features from two sub-networks. 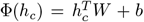, where W are the regression coefficients of size 2*D*_*L*_ ×1, and b is bias. Finally, we employ the mean square error (MSE) as the loss function to calculate the loss between the true ΔDIs and the predicted ΔDIs for our regression task, allowing us to perform end-to-end training.

## IV. Experiment

### A. Dataset

A lane-keeping driving dataset was employed in our study to explore the EEG activities associated with a sustained-attention driving task [40]. During the experiment, lane-departure events were activated randomly while participants were cruising in a car. Participants were asked to steer the car back to the original lane as soon as possible when a lane-departure event occurred. The time between the deviation onset and the response onset is the reaction time (RT), which can be used to assess drowsiness levels. Twenty-seven subjects (ages 22-28) participated in the study, contributing a total of 62 sessions (19 subjects contributed multiple sessions).

To better develop the EEG-decoding model for the drowsiness level, we first need to ensure each session accumulate sufficient data from both drowsy and alert states. We followed the session selection criterion in [22] to perform session selection. Additionally, subjects with only one session were excluded for the within-subject test, resulting in 12 subjects being included in our experiments. To establish a baseline for alert brain condition in each session, we designated the initial 10 trials as the baseline [41].

Then, we quantified drowsiness by converting RT into a drowsiness index (DI) using a previously proposed method [22]:

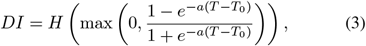

where *T* represents the RT for each trial, *T*_0_ is the alert baseline RT, *a* is a constant set to 1 *s*^−^1, and *H* is a causal moving average filter in the temporal domain.

### B. EEG data processing

The EEG signal was recorded in 30 channels with a sampling frequency of 500 Hz. First, the raw EEG data went through a bandpass filter (2-30Hz) to reduce high-frequency noise and power noise and downsampling to 250 Hz. We further applied artifact subspace reconstruction (ASR) [42] to remove the artifact in the EEG signal and set the ASR threshold as 10 times the standard deviation [22]. Finally, extract epochs that include the 3-second EEG signal prior to deviation onset. EEGLAB [43] was used to implement all data pre-processing steps.

### C. Dynamic baseline training

Based on the pairwise input, a specialized training strategy called dynamic baseline training is developed in order to enhance the adaptability and robustness of the model to variations in baseline levels. This approach involves creating pairs by randomly selecting a trial from the same session, which serves as a dynamic baseline level. The ground truth for training is established by calculating the difference in DI between these two trials (ΔDI). By introducing dynamic baseline conditions during training, dynamic baseline training forces the model to effectively adapt to varying baseline levels. Additionally, dynamic baseline training allows us to expand the dataset by controlling the number of trials used to form pairs for each trial. During the inference phase, static baseline inference is utilized. We form pairs using exclusively a fixed baseline trial. In other words, we predict the difference in DI between the current trial and a predetermined baseline trial. The resulting ΔDIs are comparable to the original DI, as they reflect the current DI after removing the baseline DI. Static baseline inference enables the evaluation of actual fluctuations in drowsiness levels during the current trial.

To assess the performance of the regression tasks, we employ two metrics: root mean square error (RMSE) and Pearson’s correlation coefficient (CC). RMSE is a commonly used metric for evaluating the disparity between the predicted ΔDIs and the actual ΔDIs, while CC measures the degree of correlation between the predicted and actual ΔDIs.

We evaluated SiamEEGNet using both within-subject and cross-subject validation. In the within-subject scheme, individual models were trained for each session, with a random 20% split for validation to assess model performance. During the testing phase, we averaged the predictions of the models trained on the same subject’s sessions. For the cross-subject scheme, we implemented leave-one-subject-out cross-validation. In the training phase, one subject from the training dataset was selected as validation data to assess model performance. The model was trained for 50 epochs using the Adam optimizer with a learning rate set to 0.001. Additionally, we employed a weight decay technique with a ratio of 0.0001 to mitigate overfitting.

### D. Selection of baseline methods

We conducted a performance comparison between SiamEEGNet and six other approaches. For the conventional drowsiness estimation methods, we utilized hand-crafted band-power features and selected support vector regression (SVR) as the classifier according to previous studies [44]–[46]. For deep learning-based methods, we chose EEGNet [11], ShallowConvNet [10], and SCCNet [12] as baseline methods to represent general single-input CNN-based approaches. Additionally, we included ESTCNN [13] and InterpretableCNN [14] as the leading DL-based EEG decoders. To ensure consistent and reliable evaluation, the experiments were repeated five times and subsequently averaged.

## V. ExperimentResult

### A. Performance comparison

First, we compared the regression performance of the proposed SiamEEGNet with 6 baseline methods. Among them, SVR used smoothed log band power features (alpha, beta, gamma and delta band log power of each channel) as input [22]. Regarding DL-based methods, we directly used the EEG signal as input and incorporated it with multi-window processing (*N*_*t*_=10) to obtain a fair comparison. Two training schemes were performed on all subjects.

Table I presents the average performance of the within-subject and cross-subject test for seven methods across 12 subjects in terms of RMSE and CC. In the within-subject test, most DL-based methods exhibit performance levels that are on par with or superior to the conventional SVR approach (RMSE=0.205, CC=0.530), with the exception of ESTCNN (RMSE=0.208, CC=0.474). Essentially, the proposed SiamEEGNet (RMSE = 0.118, CC = 0.574) can perform exceptionally well but falls slightly short of EEGNet in terms of CC. However, in the cross-subject test, the proposed model (RMSE = 0.179, CC = 0.687) outperforms all other baseline methods in terms of CC and achieves the second lowest RMSE. To assess the statistical significance of the variations among the various methods, we employed the Kruskal-Wallis test with Tukey’s HSD (Honestly Significant Difference) to perform nonparametric multiple comparisons between the methods. The results of these comparisons are presented in Table II. In the within-subject test, SiamEEGNet demonstrated significantly superior performance compared to SVR, SCCNet, and ESTCNN in CC. In the cross-subject test, SiamEEGNet statistically significantly excelled over EEGNet, SCCNet, and ShallowNet in RMSE and all baseline methods in CC. Fig. 4 presents the DI decoding results obtained by the proposed method from a sample session (S44-2). This sample DI decoding result shows that our model is capable of decoding drowsy brain dynamics and produces accurate prediction performance.

**Fig. 4:**
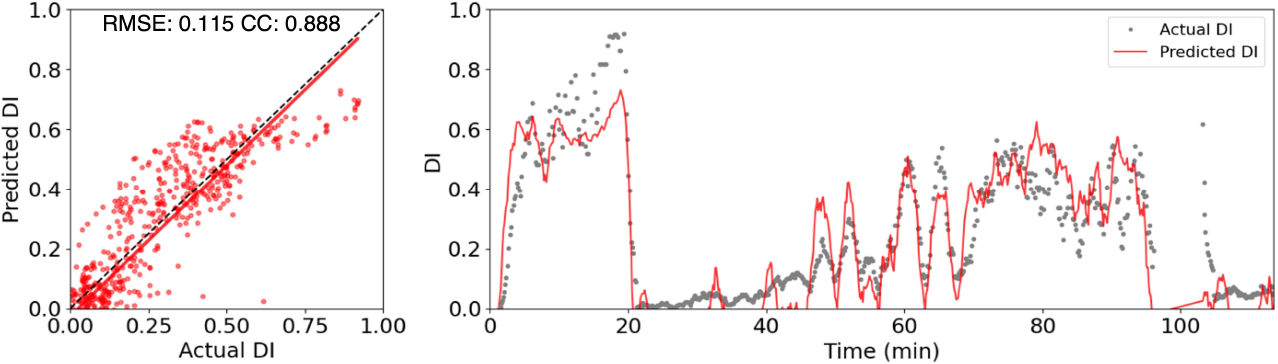
The DI decoding result of a sample session (S44-2). Left: a scatter plot of the predicted DI against the actual DI and the red line represents the linear fitting. Right: the actual DI and the temporal profile of the predicted DI.

**TABLE I:**
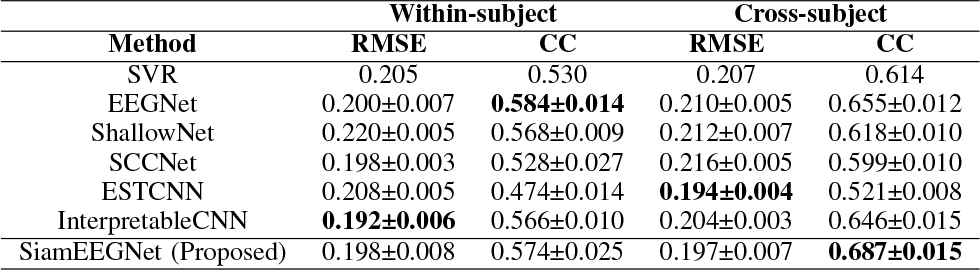
Performance comparison between the proposed SiamEEGNet and baseline leading methods in EEG-based drowsy level decoding on the overall RMSE and CC (mean±std) across all subjects.

| Method | Within-subject |  | Cross-subject |  |
| --- | --- | --- | --- | --- |
|  | RMSE | CC | RMSE | CC |
| SVR | 0.205 | 0.530 | 0.207 | 0.614 |
| EEGNet | 0.200±0.007 | **0.584±0.014** | 0.210±0.005 | 0.655±0.012 |
| ShallowNet | 0.220±0.005 | 0.568±0.009 | 0.212±0.007 | 0.618±0.010 |
| SCCNet | 0.198±0.003 | 0.528±0.027 | 0.216±0.005 | 0.599±0.010 |
| ESTCNN | 0.208±0.005 | 0.474±0.014 | **0.194±0.004** | 0.521±0.008 |
| InterpretableCNN | **0.192±0.006** | 0.566±0.010 | 0.204±0.003 | 0.646±0.015 |
| SiamEEGNet (Proposed) | 0.198±0.008 | 0.574±0.025 | 0.197±0.007 | **0.687±0.015** |

**TABLE II:** Statistical significance in the multiple comparison of within-subject and cross-subject overall performance across drowsiness detection methods. Significant differences are indicated by the p-value of the Kruskal-Wallis test with Tukey’s HSD (Honestly Significant Difference) marked in bold (p-value < 0.05).

| Statistical significance in the multiple comparison of overall performance (within-subject / cross-subject) |  |  |  |  |  |  |  |
| --- | --- | --- | --- | --- | --- | --- | --- |
|  |  | SVR | EEGNet | ShallowNet | SCCNet | ESTCNN | InterpretableCNN |
| RMSE | EEGNet | 0.7992 / 0.9529 |  |  |  |  |  |
|  | ShallowNet | 0.0724 / 0.6715 | 0.6788 / 0.9950 |  |  |  |  |
|  | SCCNet | 0.3988 / 0.1126 | 0.9933 / 0.5550 | 0.9629 / 0.8983 |  |  |  |
|  | ESTCNN | 1.0000 / **0.0072** | 0.7008 / **0.0005** | **0.0493** / **0.0001** | 0.3069 / **0.0000** |  |  |
|  | InterpretableCNN | **0.0108** / 0.9782 | 0.2273 / 0.5419 | 0.9815 / 0.2121 | 0.5894 / **0.0170** | **0.0069** / 0.0538 |  |
|  | SiamEEGNet | 0.9943 / 0.0574 | 0.9880 / **0.0053** | 0.2546 / **0.0010** | 0.7882 / **0.0000** | 0.9803 / 0.9742 | 0.0504 / 0.2931 |
| CC | EEGNet | **0.0003** / **0.0001** |  |  |  |  |  |
|  | ShallowNet | 0.0676 / 0.9969 | 0.3676 / **0.0002** |  |  |  |  |
|  | SCCNet | 1.0000 / 0.3618 | **0.0002** / **0.0000** | **0.0400** / 0.1316 |  |  |  |
|  | ESTCNN | 0.2060 / **0.0000** | **0.0000** / **0.0000** | **0.0001** / **0.0000** | 0.3067 / **0.0000** |  |  |
|  | InterpretableCNN | **0.0316** / **0.0017** | 0.5682 / 0.8487 | 0.9999 / **0.0075** | **0.0180** / **0.0000** | **0.0001** / **0.0000** |  |
|  | SiamEEGNet | **0.0051** / **0.0000** | 0.9385 / **0.0014** | 0.9296 / **0.0000** | **0.0028** / **0.0000** | **0.0000** / **0.0000** | 0.9887 / **0.0000** |

### B. Impact of feature extractors

To assess the adaptability and effectiveness of the concept of relative change in enhancing drowsiness estimation, we incorporated various EEG decoding models, not solely limited to using EEGNet as the feature extraction module. These models include five baseline DL-based models and two additional models, namely EEGTCNet [47] and MBEEGSE [48], which have shown promising performance in motor imagery classification tasks. We compared the performance with and without the proposed SiamEEGNet in the cross-subject scenario. As shown in Table III, most EEG decoding models exhibit a significant improvement when SiamEEGNet is applied, particularly in terms of CC. These results show that SiamEEGNet can be effectively integrated with various EEG decoding models, serving as a feature extraction module, and significantly elevating decoding performance, particularly in terms of CC. These results further emphasize that the concept of capturing relative changes in drowsiness levels holds universal applicability and is not confined to specific EEG decoding models.

**TABLE III:** Performance comparison between single-branch and Siamese architectures with various EEG decoding models as the backbone feature extraction module. The asterisk ‘*’ indicates a significant improvement in the overall performance obtained by using the Siamese architecture (p-value < 0.05 using the Mann-Whitney U test).

| Backbone module | #parameters within module | Single-branch architecture |  | Siamese architecture |  |
| --- | --- | --- | --- | --- | --- |
|  |  | RMSE | CC | RMSE | CC |
| EEGNet | 9513 | 0.210 | 0.655 | 0.197* | 0.687* |
| ShallowNet | 49121 | 0.212 | 0.618 | 0.209 | 0.669* |
| SCCNet | 11807 | 0.216 | 0.599 | 0.201* | 0.660* |
| EEGTCNet | 11241 | 0.209 | 0.658 | 0.200 | 0.684* |
| MBEEGSE | 8241 | 0.210 | 0.654 | 0.215 | 0.646 |
| ESTCNN | 2441093 | 0.194 | 0.521 | 0.237 | 0.566* |
| InterpretableCNN | 4625 | 0.201 | 0.646 | 0.203 | 0.647 |

### C. Impact of parameters

A series of experiments were conducted to evaluate performance under different adjustable parameter configurations. Specifically, two parameters were considered: the number of windows used in the multi-window processing and the number of dynamic baselines employed. To determine the optimal interval for achieving the best performance, various numbers of windows were tested for the multi-window processing. The changes in RMSE and CC were analyzed as a function of the number of windows, as shown in Fig. 6(a) and Fig. 6(b). The results indicate that performance initially improves with an increasing number of windows, but after 15 trials, performance reaches a plateau and even exhibited a slight decline. Based on these findings, it was determined that 10 windows would be the optimal setting for this work. In terms of time, the time span of 10 windows corresponds to approximately 90 seconds, which aligns with the filter length applied to the DIs.

Another adjustable parameter is the number of dynamic baselines used in forming input pairs. As stated previously, dynamic baseline training allows for dataset expansion by controlling the number of dynamic baselines paired with each trial. The experiments were conducted using different numbers of dynamic baselines paired with each individual trial. The results, as shown in Fig. 6(b), demonstrate consistent performance, even when the size of the dataset increased. This suggests that the size of the original training dataset is sufficient to effectively train the model with EEGNet as the feature extraction module. This can be attributed to the inherent capabilities of the model itself. EEG decoding models are typically designed to adapt to EEG datasets, which are often small-scale. Consequently, these models may lack the capacity to effectively learn from larger datasets.

### D. Ablation study

We conducted ablation studies to scrutinize the contribution of three critical components in our method, namely the Siamese architecture, dynamic baseline training, and multiwindow processing. Specifically, we removed each of these three methods separately from SiamEEGNet. Subsequently, we evaluated the regression performance of these three models in the cross-subject test. As shown in Table IV, we observed that removing the Siamese architecture resulted in a significant performance drop compared to the original SiamEEGNet. Specifically, there was a decrease of 0.013 in RMSE and 0.032 in CC. Furthermore, the use of dynamic baseline training also has a significant impact on performance. The results show a significant drop of 0.08 in RMSE and 0.031 in CC when we replaced dynamic baseline training with static baseline training. Static baseline training uses only a predefined baseline to pair with each trial during model training. Lastly, our results also reveal that multi-window processing has a significant impact on performance. When we removed multi-window processing (i.e., using only one EEG trial as input for each branch), we observed a considerable decline in decoding performance, with a reduction of 0.075 in RMSE and 0.282 in CC. To scrutinize the performance of the three variant approaches in comparison to the original SiamEEGNet in decoding the DI level, Fig. 5 displays a scatter plot of a sample session (S44-2). Although all approaches exhibit a similar trend in predicting the drowsiness index, the methods that incorporate multi-window processing display a relatively concentrated DI decoding result with less fluctuation. Additionally, dynamic baseline training demonstrates a noticeable improvement in fitting the baseline, which refers to the initial phase of a session. On the contrary, static baseline training exhibits a bias at the baseline level between actual DI and predicted DI.

**TABLE IV:** Ablation analysis of the proposed SiamEEGNet compared to variations with components excluded (\**p*-value*<* 0.05 by Mann-Whitney U test).

| Method | RMSE | CC |
| --- | --- | --- |
| SiamEEGNet (proposed) | 0.197 | 0.687 |
| w/o Siamese architecture | 0.210* | 0.655* |
| w/o Multi-window processing | 0.272* | 0.405* |
| w/o Dynamic baseline training | 0.277* | 0.656* |

**Fig. 5:**
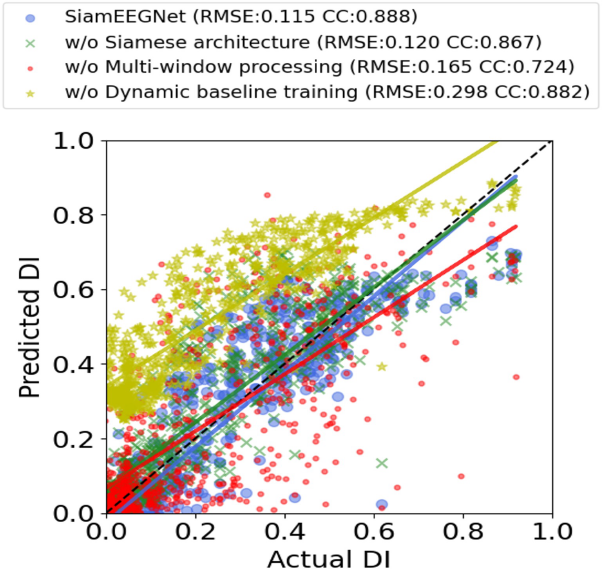
The DI decoding scatter plot with linear fitting for the ablation studies on a sample session (S44-2).

**Fig. 6:**
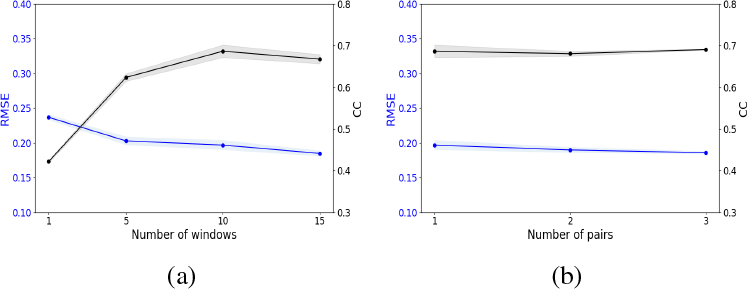
DI decoding performance (blue: RMSE; black: CC) in relation to: (a) the number of windows included in the input data for multi-window processing; (b) the number of pairs in dynamic baseline training.

## VI. Discussion

In this study, we introduce the concept of relative change, a common concept in traditional EEG analysis, to DL-based methods to improve the performance of drowsy brain activity decoding. We then developed SiamEEGNet, which enables DL-based EEG decoding models to learn the relative change between EEG trials. Our experimental results demonstrate that the proposed method outperforms all baseline methods in decoding drowsy brain activities in the cross-subject test. Compared to the result of the within-subject test, the averaged RMSE decreases by approximately 0.034 (0.206 to 0.184) while the averaged CC increases by approximately 0.113 (0.574 to 0.687). Previous studies have demonstrated that the performance of self-decoding approaches (within-subject test) is significantly affected by the inter-session variability, due to the limited availability of data from the same subjects [22]. In the cross-subject scenario, the models could access diverse data from different subjects. This factor can potentially boost the performance of models if the inter-subject variability is handled appropriately. Another factor is that the utilization of relative changes serves for enhancing the generalizability and robustness when analyzing the dynamics of the drowsy brain wave [23], [24], [49].Theses characteristics contribute to improve the performance of drowsiness detection in the cross-subject scenario.

We further conducted the ablation study to investigate the importance of learning relative changes in the decoding of drowsiness-related brain dynamics. The experiment results show that using the proposed SiamEEGNet to learn the relative change in drowsiness level significantly improves the decoding performance compared to the general single-branch EEG decoding with multi-window processing (*p* -value *<* 0.05 using the Mann-Whitney U test). These results highlight the importance of the Siamese architecture in capturing relative change. Furthermore, the way we construct input pairs for model training has a significant influence on the decoding results. We tested two different ways to form input pairs: static baseline training and dynamic baseline training. Dynamic baseline training resulted in better performance and baseline alignment in different subjects. This result shows that randomly selecting a trial as a dynamic baseline helps the models adjust to different baseline levels, which can reduce variability in the alert baseline condition between different subjects [41]. In addition, incorporating multi-window processing is another crucial factor that affects the decoding results and shows a significant improvement over not using multi-window processing. This improvement is due to the use of average latent features, which helps mitigate short-term variations and instability in EEG recordings [50] and leads to a relatively robust performance.

Through the analysis for the interpretation of the proposed model, SiamEEEGNet, we are able to uncover the underlying characteristics learned from the data. First, to explore how considering relative change can mitigate inter-subject variability and perform well in the cross-subject scenario, we performed layer-wise t-SNE [51] to visualize intermediate latent features in a 2D space. This visualization helps us to understand how latent features from different sessions evolve during the prediction process. Fig. 7 shows the layer-wise tSNE visualization of 4 example sessions (S40-1, S40-2, S44-2 and S44-3) from different layers. Initially, the latent features of the different sessions are intertwined and do not distinctly form clusters associated with the sessions. However, from the perspective of the drowsiness index, the features with higher DI tend to aggregate, while the features with lower DI are widely dispersed. Upon the application of an average pooling to latent features, sessions with lower variability (S40-1, S44-2, S44-3) exhibit a higher level of convergence. Notably, when the smoothed features are concatenated with the baseline feature of their respective sessions, features originating from the same session are anchored by their baseline and coalesce into distinct clusters. Furthermore, an increase in DI correlates with the extension of features along particular orientations relative to the baseline. By effecting predictions of the relative change for each trial in conjunction with the baseline feature, as depicted in Fig. 7(d), it is possible to align EEG trials from different subjects based on their respective baseline levels. This alignment strategy serves to mitigate the impact of inter-subject variability in baseline levels. These visualization results offer valuable insights into how the consideration of relative changes in EEG-based drowsiness detection facilitates the model’s resilience to individual difference in cross-subject scenarios.

**Fig. 7:**
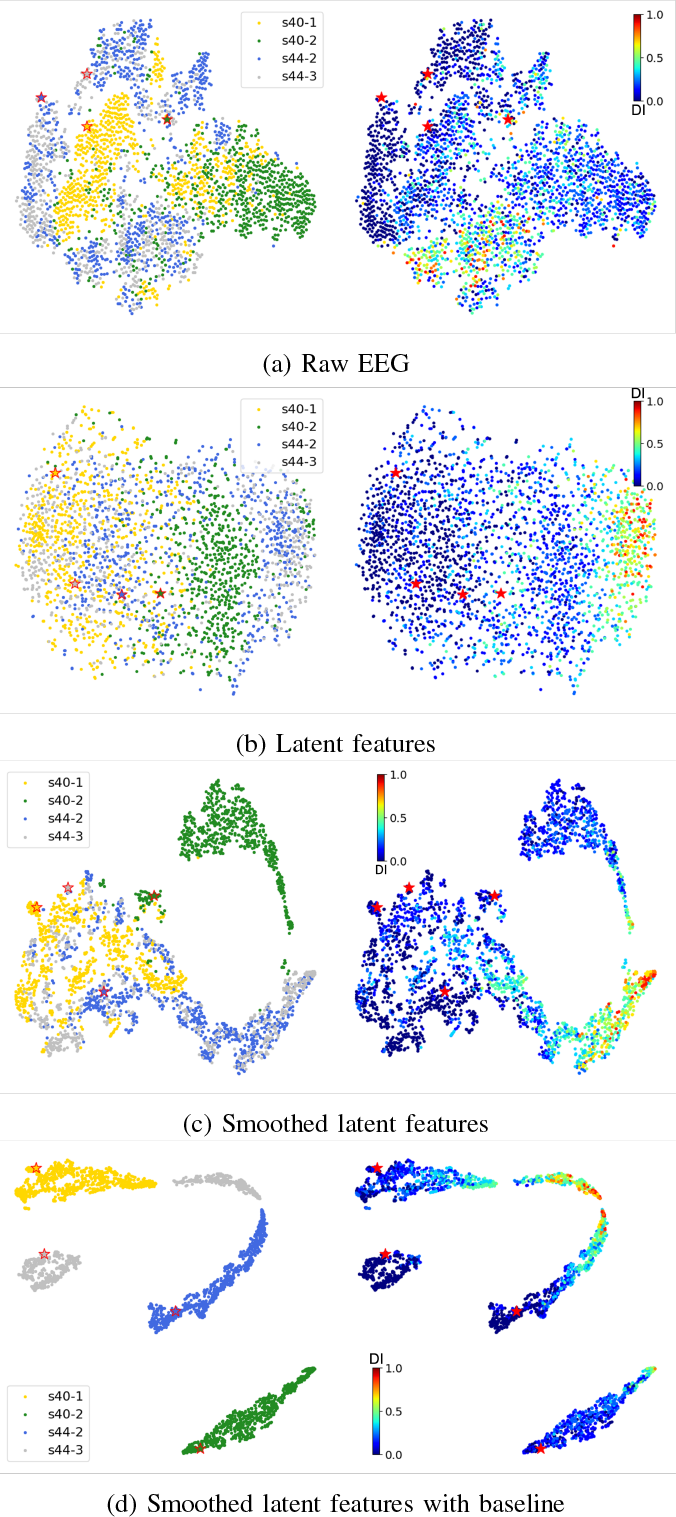
The result is a layerwise t-SNE visualization. The figures on the left are labeled by sessions, while the figures on the right are labeled by DIs. The samples marked with an asterisk are the baseline trials of the corresponding sessions. (b): output of the feature extraction module; (c): output of the average pooling layer; (c): latent feature concatenated with its baseline trial.

Then, we used saliency maps [52], [53] to investigate what our models learned from the EEG data for drowsiness detection. Saliency maps highlight important input segments and are the same size as the input (channels time × points). With this technique, we explored the important segments from two perspectives: spatial distribution and band power distribution. We analyze the spatial distribution by performing a channelwise summation to identify brain regions with high activity.

For the band power distribution, we treated each channel in the saliency map as a time sequence and calculated the power spectrum density (PSD) to discover sub-band powers of gradient response (Beta: 12-16 Hz, Alpha: 8-12 Hz, Theta: 4-8 Hz, Delta: 1-4 Hz) in each channel. We performed these analyzes on models trained with single-session data or cross-subject data. We also presented the correlation between DI and sub-band powers of the input EEG signal on the scalp to compare with active segments during model training. Fig. 8 provides evidence that the theta and delta bands exhibit a stronger correlation with DI [22], as they are also the dominant frequency bands highlighted in the saliency map. Furthermore, a higher proportion of salient components in the theta band in cross-subject models indicates that the power in the theta band is relatively universal across subjects for drowsiness estimation. This finding is also in line with previous studies showing that the theta band power spectrum has good discriminating power [41], [54]. Overall, these visualization results demonstrate that the model is capable of capturing meaningful drowsiness-related information to detect drowsiness. Additionally, the results suggest that the theta band power plays a pivotal role in the ability of our model to learn how to estimate the difference in drowsiness level. This is also consistent with previous research on EEG-based drowsiness monitoring, which has revealed that theta band power levels in the EEG signal are highly correlated with drowsiness and alertness [22], [41], [54].

**Fig. 8:**
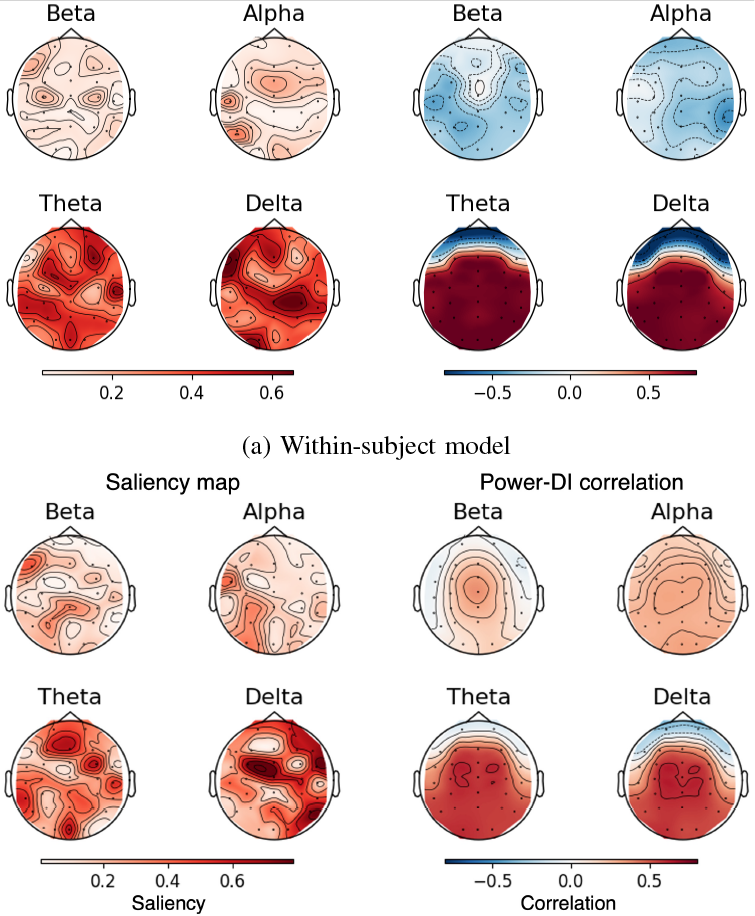
Comparison between the topoplots of saliency maps and power-DI correlation based on (a) the within-subject model (from S9-2) and (b) the cross-subject model with subject 9 (S9) as the test subject. The left side shows saliency distribution maps of four frequency bands, while the right side displays the corresponding power-DI correlation maps.

## VII. Conclusion

In this study, we introduced a Siamese neural network architecture for EEG decoding, coined as SiamEEGNet, that enables feature extraction of EEG relative change for drowsiness detection. By exploiting the characteristics of the Siamese architecture, we employed techniques of pairing EEG input to map the association between relative EEG change and relative drowsiness level difference. Furthermore, we leverage the multi-window processing and smoothing layer to address the instability and fluctuation during EEG recording, thereby enhancing the overall performance, and yield increased resilience against data variability. Moreover, the interpretation of model reveals the drowsiness-related EEG patterns and model behaviors that explain the enhanced decoding performance, which provides insights into both neuroscientific research and future EEG-based drowsiness detection. Overall, this study demonstrates the usefulness of SiamEEGNet toward highperformance practical EEG-based drowsiness detection in realworld applications.

## References

[1] Q. Maia, M. A. Grandner, J. Findley, and I. Gurubhagavatula, “Short and long sleep duration and risk of drowsy driving and the role of subjective sleep insufficiency,” Accident Analysis & Prevention, vol. 59, pp. 618–622, 2013.

[2] A. Moradi, S. S. H. Nazari, and K. Rahmani, “Sleepiness and the risk of road traffic accidents: A systematic review and meta-analysis of previous studies,” Transportation research part F: traffic psychology and behaviour, vol. 65, pp. 620–629, 2019.

[3] V. Saini and R. Saini, “Driver drowsiness detection system and techniques: a review,” International Journal of Computer Science and Information Technologies, vol. 5, no. 3, pp. 4245–4249, 2014.

[4] R. Chai, Y. Tran, G. R. Naik, T. N. Nguyen, S. H. Ling, A. Craig, and H. T. Nguyen, “Classification of eeg based-mental fatigue using principal component analysis and bayesian neural network,” in 2016 38th Annual International Conference of the IEEE Engineering in Medicine and Biology Society (EMBC). IEEE, 2016, pp. 4654–4657.

[5] S. K. Lal and A. Craig, “Driver fatigue: electroencephalography and psychological assessment,” Psychophysiology, vol. 39, no. 3, pp. 313–321, 2002.

[6] W. Klimesch, “Eeg alpha and theta oscillations reflect cognitive and memory performance: a review and analysis,” Brain research reviews, vol. 29, no. 2-3, pp. 169–195, 1999.

[7] D. D. Garrett, N. Kovacevic, A. R. McIntosh, and C. L. Grady, “The importance of being variable,” Journal of Neuroscience, vol. 31, no. 12, pp. 4496–4503, 2011.

[8] M. Rashid, N. Sulaiman, A. PP Abdul Majeed, R. M. Musa, B. S. Bari, S. Khatun et al., “Current status, challenges, and possible solutions of eeg-based brain-computer interface: a comprehensive review,” Frontiers in neurorobotics, p. 25, 2020.

[9] H. Morioka, A. Kanemura, J.-i. Hirayama, M. Shikauchi, T. Ogawa, S. Ikeda, M. Kawanabe, and S. Ishii, “Learning a common dictionary for subject-transfer decoding with resting calibration,” NeuroImage, vol. 111, pp. 167–178, 2015.

[10] R. T. Schirrmeister, J. T. Springenberg, L. D. J. Fiederer, M. Glasstetter, K. Eggensperger, M. Tangermann, F. Hutter, W. Burgard, and T. Ball, “Deep learning with convolutional neural networks for eeg decoding and visualization,” Human brain mapping, vol. 38, no. 11, pp. 5391–5420, 2017.

[11] V. J. Lawhern, A. J. Solon, N. R. Waytowich, S. M. Gordon, C. P. Hung, and B. J. Lance, “Eegnet: a compact convolutional neural network for eeg-based brain–computer interfaces,” Journal of neural engineering, vol. 15, no. 5, p. 056013, 2018.

[12] C.-S. Wei, T. Koike-Akino, and Y. Wang, “Spatial component-wise convolutional network (sccnet) for motor-imagery eeg classification,” in 2019 9th International IEEE/EMBS Conference on Neural Engineering (NER). IEEE, 2019, pp. 328–331.

[13] Z. Gao, X. Wang, Y. Yang, C. Mu, Q. Cai, W. Dang, and S. Zuo, “Eeg-based spatio–temporal convolutional neural network for driver fatigue evaluation,” IEEE transactions on neural networks and learning systems, vol. 30, no. 9, pp. 2755–2763, 2019.

[14] J. Cui, Z. Lan, O. Sourina, and W. Müller-Wittig, “Eeg-based cross-subject driver drowsiness recognition with an interpretable convolutional neural network,” IEEE Transactions on Neural Networks and Learning Systems, 2022.

[15] J.-H. Jeong, B.-W. Yu, D.-H. Lee, and S.-W. Lee, “Classification of drowsiness levels based on a deep spatio-temporal convolutional bidirectional lstm network using electroencephalography signals,” Brain sciences, vol. 9, no. 12, p. 348, 2019.

[16] D. Gao, P. Li, M. Wang, Y. Liang, S. Liu, J. Zhou, L. Wang, and Y. Zhang, “Csf-gtnet: A novel multi-dimensional feature fusion network based on convnext-gelu-bilstm for eeg-signals-enabled fatigue driving detection,” IEEE Journal of Biomedical and Health Informatics, 2023.

[17] V. Delvigne, H. Wannous, J.-P. Vandeborre, L. Ris, and T. Dutoit, “Spatio-temporal analysis of transformer based architecture for attention estimation from eeg,” in 2022 26th International Conference on Pattern Recognition (ICPR). IEEE, 2022, pp. 1076–1082.

[18] J. Wang, Y. Xu, J. Tian, H. Li, W. Jiao, Y. Sun, and G. Li, “Driving fatigue detection with three non-hair-bearing eeg channels and modified transformer model,” Entropy, vol. 24, no. 12, p. 1715, 2022.

[19] W. Zhang, F. Wang, S. Wu, Z. Xu, J. Ping, and Y. Jiang, “Partial directed coherence based graph convolutional neural networks for driving fatigue detection,” Review of Scientific Instruments, vol. 91, no. 7, p. 074713, 2020.

[20] H. Wang, L. Xu, A. Bezerianos, C. Chen, and Z. Zhang, “Linking attention-based multiscale cnn with dynamical gcn for driving fatigue detection,” IEEE Transactions on Instrumentation and Measurement, vol. 70, pp. 1–11, 2020.

[21] S. J. Luck, An introduction to the event-related potential technique. MIT press, 2014.

[22] C.-S. Wei, Y.-P. Lin, Y.-T. Wang, C.-T. Lin, and T.-P. Jung, “A subject-transfer framework for obviating inter-and intra-subject variability in eeg-based drowsiness detection,” NeuroImage, vol. 174, pp. 407–419, 2018.

[23] A. Picot, S. Charbonnier, and A. Caplier, “On-line automatic detection of driver drowsiness using a single electroencephalographic channel,” in 2008 30th Annual International Conference of the IEEE Engineering in Medicine and Biology Society. IEEE, 2008, pp. 3864–3867.

[24] G. Li, B.-L. Lee, and W.-Y. Chung, “Smartwatch-based wearable eeg system for driver drowsiness detection,” IEEE Sensors Journal, vol. 15, no. 12, pp. 7169–7180, 2015.

[25] K.-E. Ko, H.-C. Yang, and K.-B. Sim, “Emotion recognition using eeg signals with relative power values and bayesian network,” International Journal of Control, Automation and Systems, vol. 7, no. 5, p. 865, 2009.

[26] S. Markovska-Simoska and N. Pop-Jordanova, “Quantitative eeg in children and adults with attention deficit hyperactivity disorder: comparison of absolute and relative power spectra and theta/beta ratio,” Clinical EEG and neuroscience, vol. 48, no. 1, pp. 20–32, 2017.

[27] P. Golnar-Nik, S. Farashi, and M.-S. Safari, “The application of eeg power for the prediction and interpretation of consumer decision-making: A neuromarketing study,” Physiology & behavior, vol. 207, pp. 90–98, 2019.

[28] Y. Cui, Y. Xu, and D. Wu, “Eeg-based driver drowsiness estimation using feature weighted episodic training,” IEEE transactions on neural systems and rehabilitation engineering, vol. 27, no. 11, pp. 2263–2273, 2019.

[29] Y. Liu, Z. Lan, J. Cui, O. Sourina, and W. Müller-Wittig, “Inter-subject transfer learning for eeg-based mental fatigue recognition,” Advanced Engineering Informatics, vol. 46, p. 101157, 2020.

[30] J. Bromley, I. Guyon, Y. LeCun, E. Säckinger, and R. Shah, “Signature verification using a” siamese” time delay neural network,” Advances in neural information processing systems, vol. 6, 1993.

[31] A. Vaswani, N. Shazeer, N. Parmar, J. Uszkoreit, L. Jones, A. N. Gomez, Ł. Kaiser, and I. Polosukhin, “Attention is all you need,” Advances in neural information processing systems, vol. 30, 2017.

[32] D. Merkx and S. L. Frank, “Human sentence processing: Recurrence or attention?” arXiv preprint arXiv:2005.09471, 2020.

[33] H. Wang, X. Liu, J. Li, T. Xu, A. Bezerianos, Y. Sun, and F. Wan, “Driving fatigue recognition with functional connectivity based on phase synchronization,” IEEE Transactions on Cognitive and Developmental Systems, vol. 13, no. 3, pp. 668–678, 2020.

[34] Y. Taigman, M. Yang, M. Ranzato, and L. Wolf, “Deepface: Closing the gap to human-level performance in face verification,” in Proceedings of the IEEE conference on computer vision and pattern recognition, 2014, pp. 1701–1708.

[35] L. Bertinetto, J. Valmadre, J. F. Henriques, A. Vedaldi, and P. H. Torr, “Fully-convolutional siamese networks for object tracking,” in European conference on computer vision. Springer, 2016, pp. 850–865.

[36] S. Zhang, K. K. Ang, D. Zheng, Q. Hui, X. Chen, Y. Li, N. Tang, E. Chew, R. Y. Lim, and C. Guan, “Learning eeg representations with weighted convolutional siamese network: A large multi-session post-stroke rehabilitation study,” IEEE Transactions on Neural Systems and Rehabilitation Engineering, vol. 30, pp. 2824–2833, 2022.

[37] S. Shahtalebi, A. Asif, and A. Mohammadi, “Siamese neural networks for eeg-based brain-computer interfaces,” in 2020 42nd Annual International Conference of the IEEE Engineering in Medicine & Biology Society (EMBC). IEEE, 2020, pp. 442–446.

[38] Y. Li, J. Xiang, and T. Kesavadas, “Convolutional correlation analysis for enhancing the performance of ssvep-based brain-computer interface,” IEEE Transactions on Neural Systems and Rehabilitation Engineering, vol. 28, no. 12, pp. 2681–2690, 2020.

[39] X. Zhang, S. Qiu, Y. Zhang, K. Wang, Y. Wang, and H. He, “Bidirectional siamese correlation analysis method for enhancing the detection of ssveps,” Journal of Neural Engineering, vol. 19, no. 4, p. 046027, 2022.

[40] Z. Cao, C.-H. Chuang, J.-K. King, and C.-T. Lin, “Multi-channel eeg recordings during a sustained-attention driving task,” Scientific data, vol. 6, no. 1, pp. 1–8, 2019.

[41] N. R. Pal, C.-Y. Chuang, L.-W. Ko, C.-F. Chao, T.-P. Jung, S.-F. Liang, and C.-T. Lin, “Eeg-based subject-and session-independent drowsiness detection: an unsupervised approach,” EURASIP Journal on Advances in Signal Processing, vol. 2008, pp. 1–11, 2008.

[42] T. Mullen, C. Kothe, Y. M. Chi, A. Ojeda, T. Kerth, S. Makeig, G. Cauwenberghs, and T.-P. Jung, “Real-time modeling and 3d visualization of source dynamics and connectivity using wearable eeg,” in 2013 35th annual international conference of the IEEE engineering in medicine and biology society (EMBC). IEEE, 2013, pp. 2184–2187.

[43] A. Delorme and S. Makeig, “Eeglab: an open source toolbox for analysis of single-trial eeg dynamics including independent component analysis,” Journal of neuroscience methods, vol. 134, no. 1, pp. 9–21, 2004.

[44] S. K. Lal, A. Craig, P. Boord, L. Kirkup, and H. Nguyen, “Development of an algorithm for an eeg-based driver fatigue countermeasure,” Journal of safety Research, vol. 34, no. 3, pp. 321–328, 2003.

[45] C.-T. Lin, R.-C. Wu, S.-F. Liang, W.-H. Chao, Y.-J. Chen, and T.-P. Jung, “Eeg-based drowsiness estimation for safety driving using independent component analysis,” IEEE Transactions on Circuits and Systems I: Regular Papers, vol. 52, no. 12, pp. 2726–2738, 2005.

[46] C.-S. Wei, Y.-T. Wang, C.-T. Lin, and T.-P. Jung, “Toward drowsiness detection using non-hair-bearing eeg-based brain-computer interfaces,” IEEE transactions on neural systems and rehabilitation engineering, vol. 26, no. 2, pp. 400–406, 2018.

[47] T. M. Ingolfsson, M. Hersche, X. Wang, N. Kobayashi, L. Cavigelli, and L. Benini, “Eeg-tcnet: An accurate temporal convolutional network for embedded motor-imagery brain–machine interfaces,” in 2020 IEEE International Conference on Systems, Man, and Cybernetics (SMC). IEEE, 2020, pp. 2958–2965.

[48] G. A. Altuwaijri, G. Muhammad, H. Altaheri, and M. Alsulaiman, “A multi-branch convolutional neural network with squeeze-and-excitation attention blocks for eeg-based motor imagery signals classification,” Diagnostics, vol. 12, no. 4, p. 995, 2022.

[49] S. Majumder, B. Guragain, C. Wang, and N. Wilson, “On-board drowsiness detection using eeg: Current status and future prospects,” in 2019 IEEE International Conference on Electro Information Technology (EIT). IEEE, 2019, pp. 483–490.

[50] T.-P. Jung, S. Makeig, M. Stensmo, and T. J. Sejnowski, “Estimating alertness from the eeg power spectrum,” IEEE transactions on biomedical engineering, vol. 44, no. 1, pp. 60–69, 1997.

[51] L. Van der Maaten and G. Hinton, “Visualizing data using t-sne.” Journal of machine learning research, vol. 9, no. 11, 2008.

[52] K. Simonyan, A. Vedaldi, and A. Zisserman, “Deep inside convolutional networks: Visualising image classification models and saliency maps,” arXiv preprint arXiv:1312.6034, 2013.

[53] J. T. Springenberg, A. Dosovitskiy, T. Brox, and M. Riedmiller, “Striving for simplicity: The all convolutional net,” arXiv preprint arXiv:1412.6806, 2014.

[54] B. T. Jap, S. Lal, P. Fischer, and E. Bekiaris, “Using eeg spectral components to assess algorithms for detecting fatigue,” Expert Systems with Applications, vol. 36, no. 2, pp. 2352–2359, 2009.

